# Matrin3 regulates cell proliferation and spindle dynamics by regulating CDC14B alternative splicing

**DOI:** 10.1101/2022.06.08.495349

**Authors:** Bruna R. Muys, Roshan L. Shrestha, Dimitrios G. Anastasakis, Lorinc Pongor, Xiao Ling Li, Ioannis Grammatikakis, Ahsan Polash, Curtis C Harris, Mirit I. Aladjem, Munira A. Basrai, Markus Hafner, Ashish Lal

## Abstract

Matrin3 is an RNA-binding protein that affects diverse RNA-related processes, including mRNA splicing. While Matrin3 has been intensively studied in neurodegenerative diseases, its function in cancer remains unclear. Here, we discovered Matrin3-mediated regulation of mitotic spindle dynamics in colorectal cancer (CRC) cells. We identified bound and regulated Matrin3-target RNAs transcriptome-wide in CRC cells and found that Matrin3 broadly modulates mRNA splicing patterns. Among the top Matrin3 targets, we focused on *CDC14B* and found that Matrin3 loss increased inclusion of an exon containing a premature termination codon into the *CDC14B* transcript and simultaneous down-regulation of the standard *CDC14B* transcript. Selective knockdown of the *CDC14B* standard transcript phenocopied Matrin3 knockdown and exhibited reduced proliferation and defects in mitotic spindle formation, suggesting that *CDC14B* is a key downstream effector of Matrin3. Collectively, these data reveal an important role for the *Matrin3/CDC14B* axis in control of CRC cell proliferation and mitotic spindle formation.

## Introduction

Matrin3 (MATR3) is a nucleic acid binding protein^1,2^ conserved in vertebrates that is the major component of the nuclear matrix, a highly structured residual framework mainly composed of lamins and ribonucleoproteins (RNPs)^3,4^. *In vitro*, Matrin3 can bind to DNA with its two zinc-fingers (ZF) motifs and to RNA through two tandem RNA recognition motifs (RRM)^5^. Evidence suggests that by binding to DNA, Matrin3 can attach to specific chromatin structural elements termed matrix or scaffold attachment region (MAR/SAR)^6^. Consistently, Matrin3 was recently shown to regulate chromatin organization^7,8^. Nevertheless, most studies to date focused on the role of Matrin3 as an RBP regulating many diverse - not necessarily related - RNA processes, including transcription^9,10^; splicing^11–13^, RNA stability^14^, nuclear export^15^ and nuclear retention of hyper-edited RNAs^16^. We found that after DNA damage in human colorectal cancer (CRC) cells, the long non-coding RNA (lncRNA) *PINCR* directly binds and recruits Matrin3 to enhancers of p53 target genes that depend on *PINCR* for p53-mediated induction of a subset of p53 targets (BTG2, GPX1 and RRM2B)^17^. Moreover, during myogenesis, Matrin3 is required for paraspeckle formation, likely by controlling adenosine to inosine (A-to-I) RNA editing of Ctn RNA^18^.

Global knockout of Matrin3 is embryonic lethal^19^ indicating that it is required for normal development. Furthermore, conditional deletion in the neuronal lineage showed that Matrin3 is essential for maintaining neuronal survival^20^. Even though Matrin3 appears to be essential *in vivo*, disease-associated mutations were primarily found in neurodegenerative disorders, e.g., in familial Amyotrophic Lateral Sclerosis (ALS) and frontotemporal dementia (FTD)^21^. The impaired formation of dynamic nuclear condensates due to ALS-associated mutations in Matrin3 possibly contributes to its pathogenic mechanism^22^. There is conflicting evidence for a role of Matrin3 in cancer with some studies indicating a potential role of Matrin3 as an oncogene and other studies suggesting a tumor suppressive function^23–25^. Although many different Matrin3 RNA targets were catalogued^11–13^, it remains unclear which of these specific targets mediate Matrin3 function and contribute to the observed phenotypes of Matrin3 in different cells and tissues.

Here, we expanded our findings focusing on the role of Matrin3 protein in CRC. We found that Matrin3 is overexpressed in CRC and provides growth advantage in CRC cell lines. To understand the molecular mechanism(s) by which Matrin3 mediates its effects, we performed PAR-CLIP (Photoactivatable Ribonucleoside-enhanced CrossLinking and Immunoprecipitation), a UV-crosslinking based technique that identifies RNAs directly interacting with RNA-binding proteins (RBPs) at nucleotide resolution on a transcriptome-wide scale^26,27^. We found that in HCT116 cells (CRC), Matrin3 binds to thousands of pre-mRNAs at pyrimidine-rich sequence elements repressing inclusion of nearby exons. For functional analysis, we focused on *CDC14B*, which had one of the highest changes in splicing patterns upon Matrin3 depletion. *CDC14B* encodes a protein tyrosine phosphatase that has roles in DNA damage and repair^28,29^, ciliogenesis^30^ and early-onset aging^31^. We show that Matrin3 depletion resulted in decreased expression of the most abundant CDC14B variant and CDC14B knockdown phenocopied the reduced cell proliferation and mitotic defects of Matrin3 knockdown. Collectively, these data reveal a growth-promoting function of Matrin3 in CRC cells that is - at least partially - mediated through splicing changes of CDC14B.

## Results

### Matrin3 binds to pyrimidine-rich intronic sites in nascent transcripts encoding proteins that regulate the mitotic spindle

Considering the conflicting evidence for a potential tumor suppressive or oncogenic function of Matrin3, we first determined the expression profile of Matrin3 mRNA in a panel of tumors compared to normal tissue deposited in the Oncomine database (https://www.oncomine.org/). We found that Matrin3 mRNA is significantly overexpressed in tumor tissues (≥2-fold and p<0.0001, Extended Data Fig. 1A), most frequently in CRC. We validated this finding by analyzing data from Colon Adenocarcinoma (COAD) available from the TSVdb database that collects TCGA data^32^, as well as from a cohort of CRC patients from the University of Maryland Medical Center (UMMC). In both datasets Matrin3 mRNA was significantly upregulated in tumors compared to normal tissues, suggesting a potential oncogenic function for Matrin3 in CRC (Fig.1A and Extended Data Fig. 1B respectively). Consistent with these data, Matrin3 knockdown using a pool of 4 Matrin3 siRNAs (Extended Data Fig. 1C-1D) resulted in decreased proliferation and clonogenicity in two CRC cell lines (HCT116 and LS180) (Fig. 1B and 1C and Extended Data Fig. 1E-F).

**Fig. 1.**
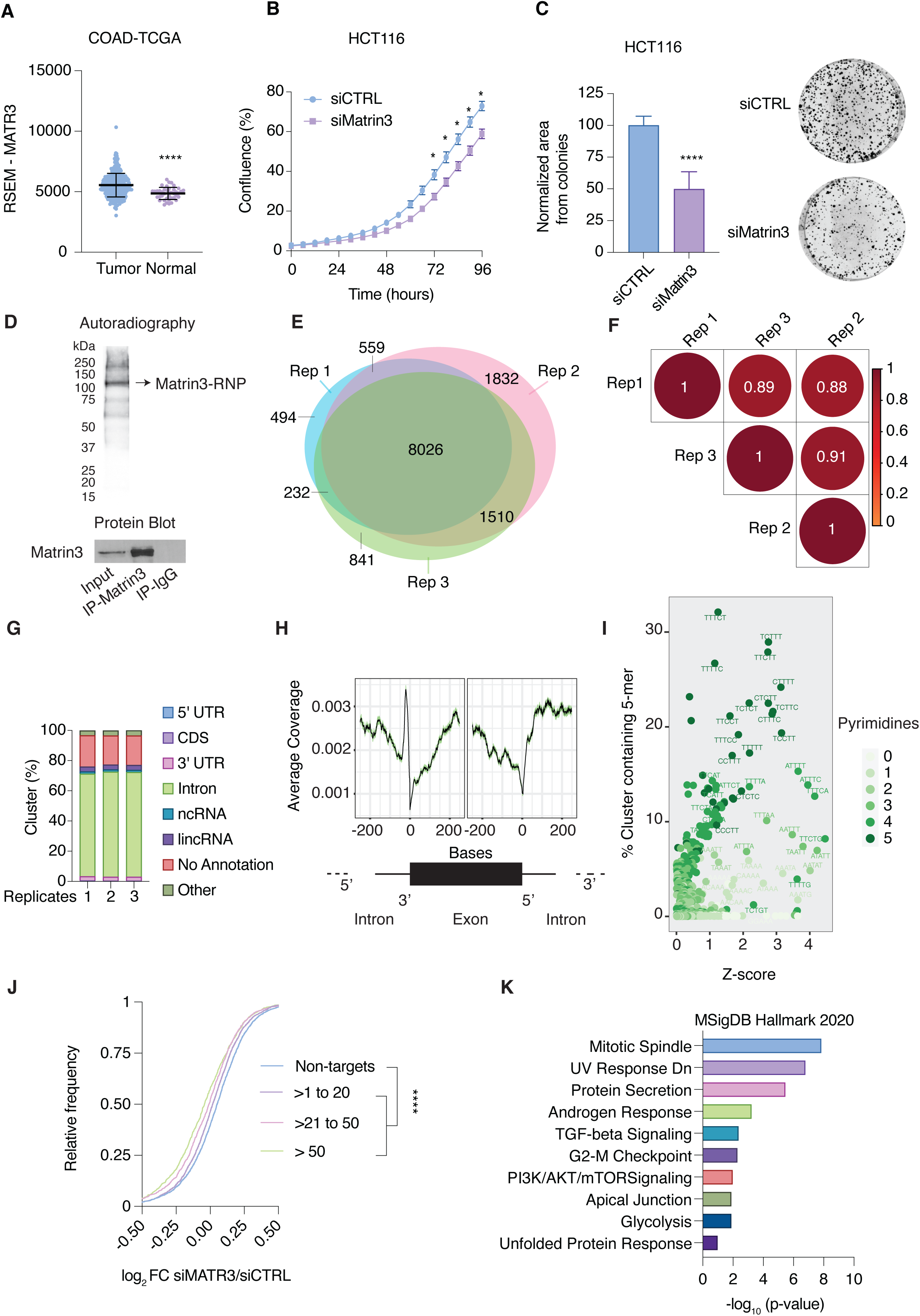
The RBP Matrin3 is a potential oncogene in colorectal cancer and binds pre-mRNAs to affect their expression. **A.** Matrin3 RNA expression in colorectal cancer (TCGA COAD) from the TSVdb database compared to respective normal samples. Unpaired, two-sided t-test; Tumor, N = 287; Normal, N = 41. Error bars, SD, ****p<0.0001. **B.** Cell proliferation assay after siRNA-knockdown of Matrin3 in HCT116 cells. Two-way ANOVA test; N = 3; Error bars = SEM; F = 13.3, DF = 1 and *p<0.05. **C.** Colony formation in HCT116 cells after Matrin3 knockdown. Unpaired, two-sided t-test; N = 3; Error bars = SD and ****p<0.0001. **D.** Autoradiography of the P32-labeled Matrin3-RNP after immunoprecipitation (IP). Band migrating at expected size of the Matrin3-RNP is indicated. (Bottom panel) Matrin3 immunoblot of Input, Matrin3-IP, and IgG-IP. **E.** Intersection of Matrin3 target transcripts identified in the three biological PAR-CLIP replicates. **F.** Spearman’s correlation matrix of crosslinked sequence read frequency (measured by number of T-to-C mutations) on target transcripts for the PAR-CLIP replicates. **G.** Average distribution of binding sites from Matrin3-bound RNAs across different annotation categories. **H.** Metagene analysis of Matrin3 binding sites around the intron-exon boundary. **I.** Scatter plot of Z-scores (x-axis) and frequency of occurrence (y-axis) of all possible 5-mers in Matrin3 PAR-CLIP binding sites. Shades of green indicate the number of pyrimidines. **J.** Cumulative distribution of gene abundance change (log_2_) determined by RNA-seq after Matrin3 knockdown and binned by normalized cross-linked reads per million (NXPM). Two-tailed Komogorov-Smirnov test; Non-targets, N = 2,579; [1 to 20 NXPM] N = 2,701; [21-50 NXPM] N = 900; [>50 NXPM] = 1,015 and ****p<0.0001. **K.** Plot of p-values of the enriched GSEA-Molecular Signature Database (MSidDB) terms constructed from top Matrin3 targets.

Matrin3 is a well-characterized RNA-binding protein (RBP) and we hypothesized that its effect on cell proliferation may be mediated by a set of specific RNA targets. Therefore, we mapped Matrin3 binding sites on RNAs on a transcriptome-wide scale and at nucleotide resolution in HCT116 cells using 4-thiouridine (4SU) PAR-CLIP^26,27^. For PAR-CLIP we immunoprecipitated (IP) endogenous, UV-crosslinked Matrin3 RNP (ribonucleoprotein) with a specific antibody. Autoradiography of crosslinked, ribonuclease-treated Matrin3 IP fractionated on denaturing protein gels revealed a prominent band migrating at the expected size of ∼125 kDa, corresponding to the Matrin3-RNP (Fig. 1D).

We recovered the RNA fragments of Matrin3-RNPs from three biological replicates and converted them into small RNA cDNA libraries for next-generation sequencing. Next, we determined clusters of overlapping reads that harbor characteristic T-to-C conversions diagnostic of 4SU-crosslinking events at higher frequencies than expected by chance^33^. For the three samples we found between 99,087 and 168,992 binding sites (Extended Data Table 1). These binding sites distributed on a set of more than 8,000 shared genes, suggesting that a significant proportion of expressed transcripts in a cell are bound by Matrin3 (Fig. 1E). Overall, the biological replicates showed excellent correlation with R^2^ of ∼0.9 (Fig. 1F). Approximately 70% of Matrin3 binding sites were found on intronic regions (Fig. 1G), indicating that Matrin3 was interacting with nascent transcripts, consistent with its nuclear localization and previous CLIP-based studies^11–13^. Some of these previous studies found a role for Matrin3 in alternative splicing, and indeed, in our data, Matrin3 bound broadly across nascent transcripts with a pronounced enrichment near the 3’ splice site (Fig. 1H), most likely the polypyrimidine tract required for splicing, considering that Matrin3 binding sites were enriched for 5-mers containing pyrimidines (Fig. 1I).

To investigate the gene regulatory roles of Matrin3, we next performed RNA-seq from HCT116 cells transfected with Matrin3 siRNAs or a negative control siRNA. Depletion of endogenous Matrin3 led to a statistically significant, albeit small decrease in target mRNA levels. The magnitude of this effect was dependent on the overall strength of binding, i.e., the number of Matrin3 binding sites per target mRNA, or the number of crosslinked reads per target mRNA normalized by overall mRNA abundance (Fig. 1J and Extended Data Table 2). We previously found that both metrics correlated well with the occupancy of an RBP on its target^34^.

We next asked which basic architectural^35^ features differentiated Matrin3 targets from non-targets. We found that Matrin3 targets were typically encoded on longer loci than non-targets. As one would expect from an RBP binding nascent transcripts, Matrin3 PAR-CLIP binding site numbers correlated with transcript length (Extended Data Fig. 2A), number of exons (Extended Data Fig. 2B), exon length (Extended Data Fig. 2C), or intron length (Extended Data Fig. 2D). Furthermore, top Matrin3 targets had lower GC content compared to weaker targets or non-targets (Extended Data Fig. 2E), consistent with the preference of RRM domains binding unstructured RNA.

To identify molecular pathways regulated by Matrin3, we performed Gene Set Enrichment Analysis (GSEA) using the GSEA-Molecular Signature Database (MSidDB). Top Matrin3 targets (with more than 50 binding sites) were enriched for genes related to the mitotic spindle (Fig. 1K), a structure important for mitosis. Thus, the PAR-CLIP interactome complemented our cell-based findings that Matrin3 knockdown resulted in reduced proliferation and clonogenicity of CRC cell lines.

### Matrin3 depletion leads to inclusion of exons proximal to its binding sites

Matrin3 has been previously characterized as a regulator of alternative splicing^11–13^. Therefore, we analyzed alternative splicing (AS) patterns in our polyA-selected RNA-seq data from HCT116 cells after Matrin3 knockdown. Using a false discovery rate (FDR) cutoff of <0.05 and ΔPSI ≥ 10% (Percent Spliced In), we found that most AS events involved cassette exons or skipped exons (SE) (Fig. 2A). Of those events, >70% were found in Matrin3 bound RNAs, suggesting a direct effect of Matrin3. As reported previously in other cell types^11,13^, Matrin3 acted mainly as a splicing repressor with 66% of the SE events in Matrin3 targets resulting from exon inclusion after Matrin3 depletion (Fig. 2B).

**Fig. 2.**
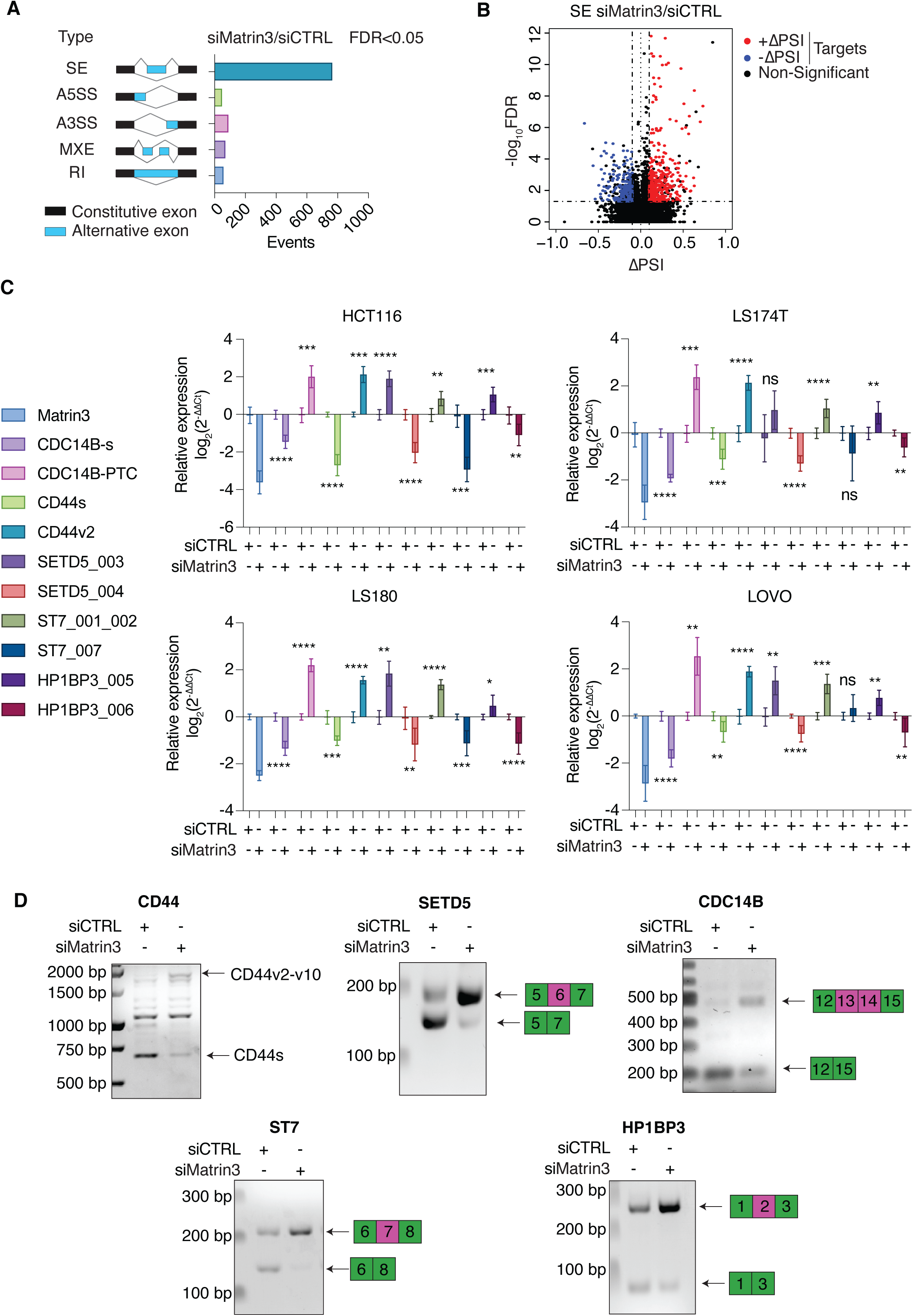
Matrin3 is a splicing suppressor and its knockdown results in widespread exon in target mRNAs. **A.** Bar plot categorizing the number of different alternative splicing events after Matrin3 knockdown in HCT116 cells (FDR <0.05 and ΔPSI ≥ 10%). **B.** Volcano plot of FDR (- log_10_) versus ΔPSI of exon skipping (SE) events upon Matrin3 silencing. Black dots, SE events with FDR >0.05; blue and red dots, exon exclusion and inclusion events, respectively with FDR <0.05 in Matrin3 target mRNAs. **C.** RT-qPCR quantification of a panel of mRNA; either full-length mRNA, or inclusion levels of indicated exons, were measured, and normalized by GAPDH after knockdown of Matrin3 in four different CRC cell lines (HCT116, LS174T, LS180, and LOVO). Unpaired, two-sided t-test; N = 3. Error bars = SD, *p<0.05, ****p<0.0001 and ns: non-significant. **D.** Representative images (N = 3) of semi-quantitative RT-PCR upon Matrin3 silencing in HCT116 cells with primers designed around alternatively spliced exon(s) of genes showed in panel C.

We next performed RT-qPCR to validate some of these AS events on transcripts with high ΔPSI in HCT116 and three other CRC cell lines (LS174T, LS180, and LOVO) after Matrin3 knockdown to expand the relevance of our results beyond HCT116. We analyzed specific exon inclusion events in the *CDC14B*, *CD44*, *SETD5*, and *HP1BP3* pre-mRNAs and were able to validate the RNA-seq results in all cases (Fig. 2C, Extended Data Fig. 3). Note that for the *ST7* pre-mRNA, we only validated the AS in the HCT116 and LS180 cell lines (Fig. 2C, Extended Data Fig. 3), still consistent with our initial observation from RNA-seq. Finally, we used semi-quantitative RT-PCR and agarose gel electrophoresis to further validate the AS event on these pre-mRNAs in HCT116 cells (Fig. 2D). For CDC14B, we named the variants as CDC14B-s and CDC14B-PTC, as described below. Taken together our results indicate that Matrin3 binding close to 3’ splice sites suppresses alternative splicing and its loss results in mRNA misprocessing that manifest in inclusion of otherwise excluded exons.

### CDC14B mediates the pro-proliferative effects of Matrin3

Among the genes that were significantly bound and changing exon inclusion patterns after silencing of Matrin3, we chose to focus on *CDC14B* for functional analysis, as it was among the genes with most significantly changed ΔPSI upon Matrin3 knockdown. We hypothesized that the significant reduction in cell proliferation after Matrin3 knockdown is mediated, at least in part, by missplicing of *CDC14B*. *CDC14B* encodes a protein tyrosine phosphatase that is similar to *Saccharomyces cerevisiae Cdc14*, and regulates interphase nuclear architecture, mitotic spindle assembly, and M phase exit^36^. We found that in all cell lines tested, Matrin3 knockdown resulted in ∼60% reduction of the most abundant CDC14B variant found in HCT116 cells (CDC14B-003 or ENST00000375241), that we refer to as “standard variant” or “CDC14B-s”. This decrease was accompanied by increased inclusion of two exons, which are only annotated in CDC14B-008 (ENST00000481149), a potential processed transcript variant that does not encode a protein. Interestingly, using Iso-seq from HCT116 cells (Extended Data Table 4), we could detect CDC14B-s as the only coding transcript in these cells implying that this the major variant expressed. We refer to the longer transcript variant as “PTC variant” or “CDC14B-PTC” (Fig. 3A) because exon 13 introduces a premature termination codon (PTC), which makes this transcript a potential substrate for non-sense mediated decay (NMD). Immunoblotting confirmed the decrease in total CDC14B protein levels after knockdown of Matrin3 (Fig. 3B). Knockdown of CDC14B-s was also validated by RT-qPCR and immunoblotting using an siRNA targeting the CDC14B-s transcript (Fig. 3B and 3C).

**Fig. 3.**
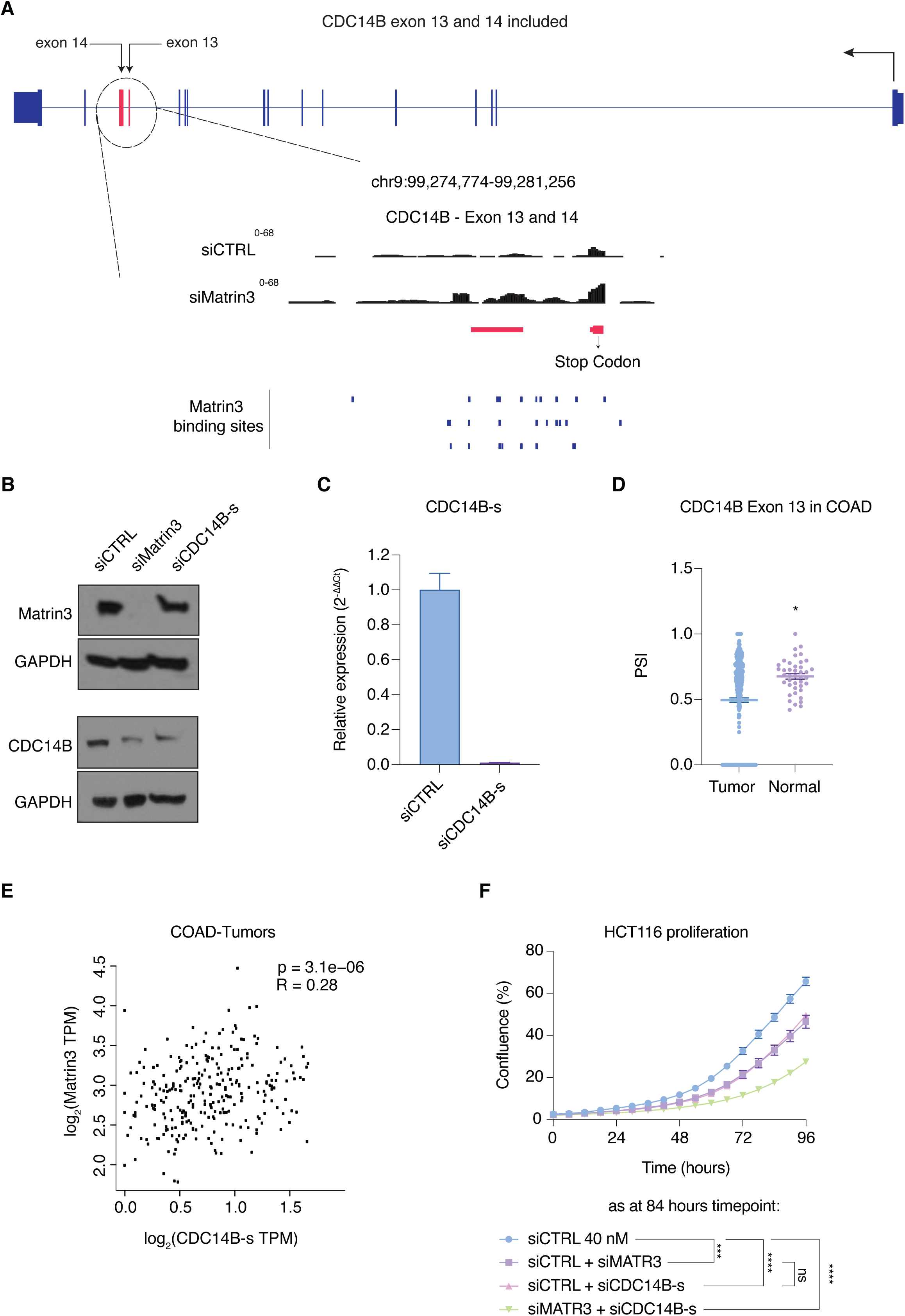
Alternative splicing of CDC14B partially mediates the Matrin3 phenotype in CRC. **A.** Genome browser image of the CDC14B-s with exons 13 and 14 included. The focused region shows tracks of sample treated with siCTRL and siMatrin3 knockdown. Lower tracks show Matrin3 RNA binding sites in the region. **B.** Immunoblots from HCT116 cell lysates after silencing of Matrin3 or CDC14B-s antibodies against Matrin3 or CDC14B, and GAPDH as control. **C.** RT-qPCR quantifying CDC14B-s variant expression after knockdown of CDC14B-s using siRNA. Error bars = SD. **D.** PSI of CDC14B exon 13 in COAD and respective normal tissues deposited in the TCGA SpliceSeq database. Unpaired, two-sided t-test; N= 3. N Tumor = 458 and N Normal = 41. Error bars = SEM, *p<0.05. **E.** Spearman’s correlation of Matrin3 and CDC14B-s expression in TCGA COAD tumor samples deposited in the Gepia2 database (http://gepia2.cancer-pku.cn/). **F.** Cell proliferation assay with Incucyte® after Matrin3 and/or CDC14B-s knockdown in HCT116 cells. Two-way ANOVA test; N = 3; Error bars = SEM; F = 48.52 and DF = 3. ***p<0.001, ****p<0.0001 and ns: non-significant.

Interestingly, in TCGA data for COAD, exon 13 is preferentially included in normal tissues compared to tumors (Fig. 3D). We also observed a modest but significant positive correlation between Matrin3 and CDC14B-s RNA expression in COAD samples (Fig. 3E). Because we found that Matrin3 loss altered cell proliferation (Fig. 1B), we next determined if it would also be affected by knockdown of CDC14B-s. Knockdown of Matrin3 or CDC14B-s in HCT116 cells resulted in a similar decrease of cell proliferation compared to siCTRL. Concurrent knockdown of Matrin3 and CDC14B-s had an additive effect on cell proliferation (Fig. 3F). Taken together, our data suggest that CDC14B splicing regulation may contribute to the decreased proliferation observed upon Matrin3 knockdown.

### Matrin3 and CDC14B regulate microtubule dynamics and orientation of the mitotic spindle

CDC14 is a conserved protein in eukaryotes. The crucial role of CDC14 in mitosis in budding yeast^37^ and the influence of Matrin3 on CDC14B expression, prompted us to examine if Matrin3 plays a role in maintaining mitotic spindle architecture. We therefore examined the morphology of mitotic spindles, microtubule dynamics and spindle orientation upon depletion of Matrin3 or CDC14B-s. To visualize the morphology of the mitotic spindle, cells were immunostained with alpha-tubulin for spindle microtubules and Nuf2 for an outer kinetochore marker. We found that in metaphase, mitotic spindles were elongated and slender in Matrin3 or CDC14B-s depleted cells compared to control cells (Fig. 4A, left panel). Next, we measured the distance between two spindle poles across the orientation of the mitotic spindle in a 2D plane (in the same z-stack) relative to the cell size. This analysis revealed that mitotic spindles were significantly elongated in cells depleted for Matrin3 or CDC14B-s (Fig. 4A, right panel). We reasoned that the elongated spindles could be caused by altered microtubule dynamics with more polymerizing microtubules present as seen in cells with more stabilized microtubules^38^. Therefore, we examined the density of EB1 comets that mark polymerizing microtubules in interphase cells upon Matrin3 or CDC14B-s depletion. Qualitative analysis showed that in interphase cells, the density of EB1 comets was higher upon Matrin3 or CDC14B-s depletion compared to control (Fig. 4B), suggesting that density of polymerizing microtubules is higher. In mitotic cells, we used EB1 signal intensity to measure the density of polymerizing microtubules, as EB1 comet analysis was not feasible due to the highly dynamic nature of microtubules in mitosis. EB1 intensity measured by imaging and pseudocoloring EB1 signal showed brighter comets (yellow) when compared to less bright comets (red) in Matrin3 or CDC14B-s depleted cells (Fig. 4C). These results confirm that the microtubules were highly polymerized in Matrin3 or CDC14B-s depleted cells.

**Fig. 4.**
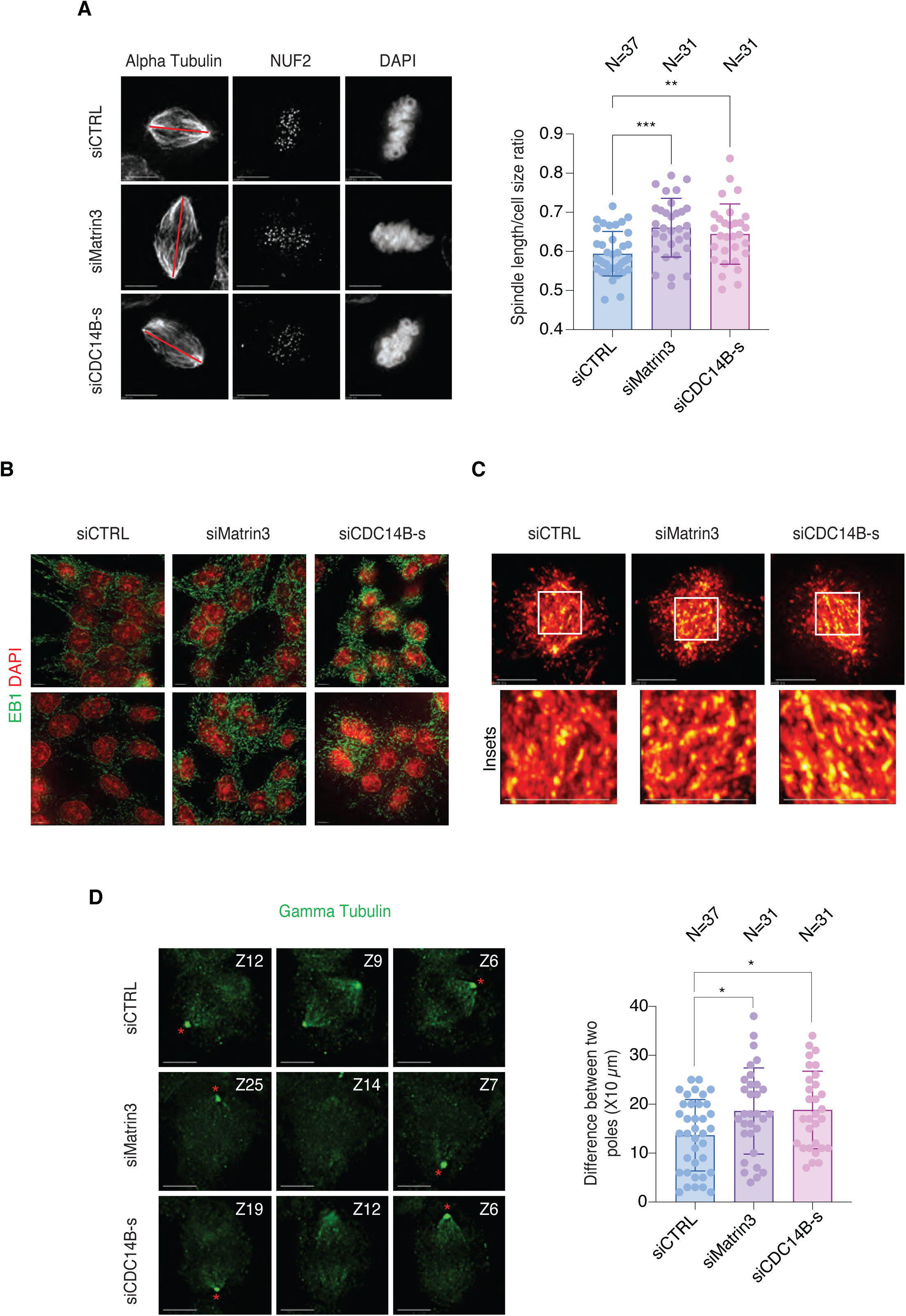
Matrin3 and CDC14B-s affect microtubule dynamics and spindle orientation. **A.** Representative immunofluorescence images of cells in metaphase stained for alpha tubulin (spindle microtubules) or Nuf2 (kinetochores) (left panel) after treatment with Matrin3 or control siRNAs. Nuclei were stained with DAPI, Scale bar: 5 µm; red bars indicate length of mitotic spindles. (Right panel) Bar graph of ratio between spindle length and cell size were calculated. “N” denotes number of cells analyzed. One-way ANOVA test; Error bars = SD; F = 8.58 and DF = 2, **p<0.01 and ***p<0.001. **B.** Representative immunofluorescence images showing EB1 comet density in interphase HCT116 cells treated with siRNAs as indicated and stained for EB1 and DAPI. Scale bar: 5 µm. **C.** Same as in B, except cells were only stained for EB1 and images pseudocolored to show highly intensified comets as yellow and less intensified comets as red. Scale bars: main image, 5 µm; insets, 1 µm. **D.** Representative metaphase images showing mitotic spindle orientation as depicted by planes of two spindle poles denoted with red asterisks in HCT116 cells treated with the indicated siRNAs and stained for gamma-tubulin (left panel). Cells were imaged with multiple 10-µm z-stacks, and number of z-stacks between planes of two poles were measured in each siRNA condition and plotted as a bar plot (right panel). “N” denotes number of cells analyzed from two independent experiments. One-way ANOVA test; Error bars = SD; F = 4.65 and DF = 2. Scale bar: 5 µm and *p<0.05.

We next examined the consequence of Matrin3 or CDC14B-s depletion on the orientation of the mitotic spindle. Previous studies have shown that during symmetric cell division, epithelial cells often orient their mitotic spindles parallel to the plane of attachment and this is regulated by several factors, including microtubule dynamics^39^. The orientation of the mitotic spindles was examined by immunostaining with gamma tubulin that marks the two ends of the mitotic spindle. Matrin3 or CDC14B-s depleted cells showed misorientation of mitotic spindles, as we did not see two poles of a mitotic spindle on the same optical plane in most of these cells (Fig. 4D, left panel). For quantitative analysis we counted z-stacks between planes of two poles (from the plane where one pole is clear to another plane where another pole is clear). Cells depleted for Matrin3, or CDC14B-s had higher z-stack counts compared to control (Fig. 4D, right panel), indicating that the two poles of a mitotic spindle in these cells are not in the sample plane supporting our conclusion for misorientation of mitotic spindles in these cells.

Taken together, as summarized in our model (Fig. 5), our data shows that by regulating the abundance of CDC14B-s, Matrin3 contributes to maintenance of microtubules dynamics, spindle morphology, and proper mitotic spindle orientation and suggest that defects in these processes may contribute to the reduced proliferation of cells after depletion of Matrin3.

**Fig. 5.**
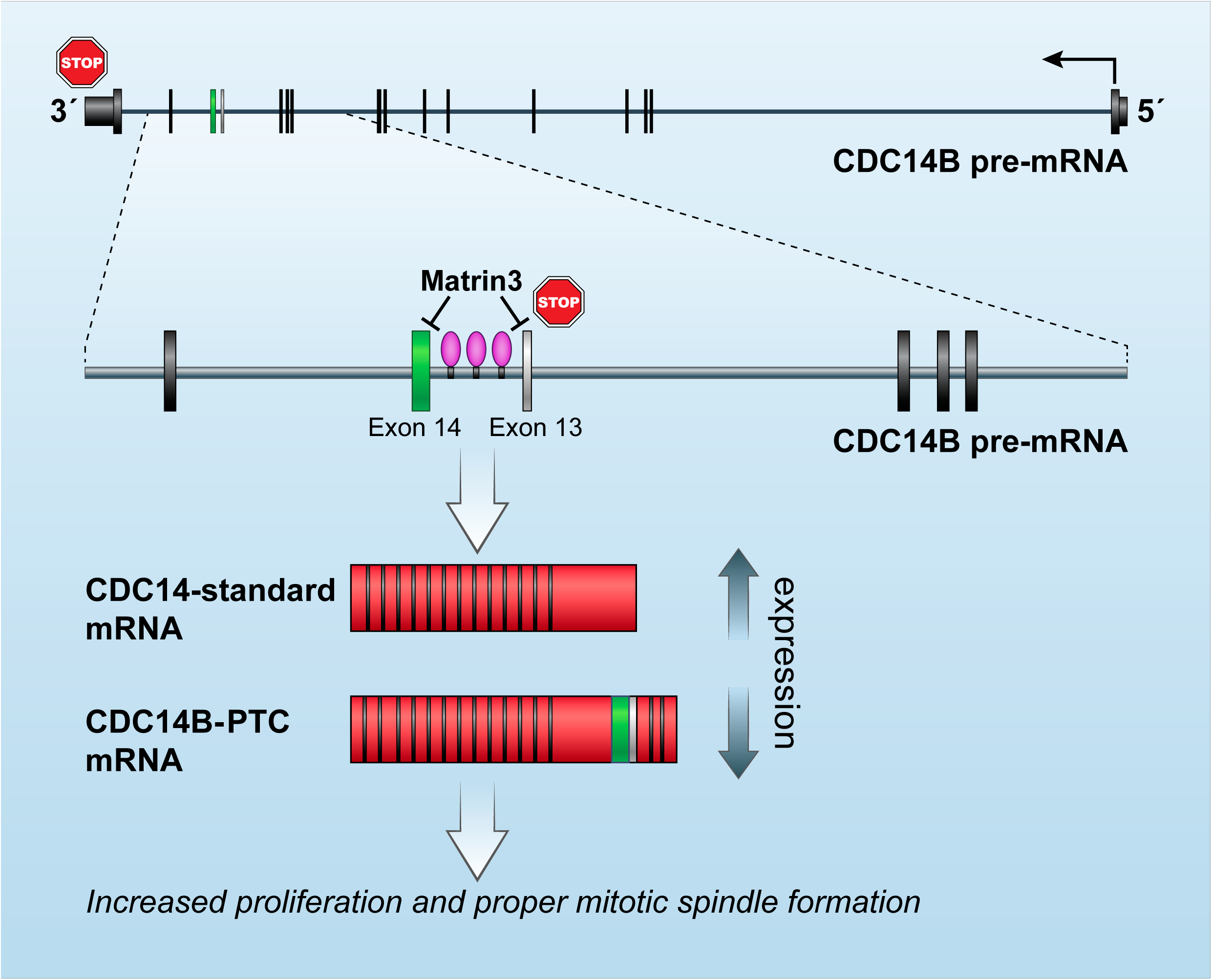
Model: Matrin3 inhibits CDC14B alternative splicing to promote proliferation and spindle dynamics in CRC cells.

## Discussion

Matrin3 is typically studied in the context of neurodegenerative disorders, like ALS^40^, with only a handful of reports documenting its involvement in cancer. Here, we demonstrated that Matrin3 is upregulated in CRC tumors and that its knockdown results in growth defects in CRC cells. We identified the bound and regulated Matrin3 target RNAs transcriptome-wide and discovered that Matrin3 promotes CRC growth by suppressing CDC14B splicing and altering microtubule dynamics.

Although several Matrin3 CLIP experiments were reported previously^11–13^, most of these reports focused on the relationship of the binding sites of Matrin3 and PTBP1, another splicing regulator. By integrating RNA-seq and PAR-CLIP data, we corroborated that Matrin3 acts mainly as a splicing repressor^11,13^. We do not necessarily interpret this as evidence for Matrin3 being a regulatory molecule, but rather as being required for proper mRNA processing. Consistently, we found that Matrin3 coats its best targets across the entire pre-mRNA (exons and introns) and loss of Matrin3 results in reduced target mRNA abundance, likely by nuclear degradation pathways dedicated to sensing misprocessed and damaged RNA^41^. Nevertheless, many of the exon inclusion events upon Matrin3 depletion will still result in alternatively spliced transcripts. Interestingly, Matrin3 loss resulted in similar mRNA expression patterns independent of the cell line, as we were able to validate alternatively spliced transcripts across multiple types of CRC cell lines, indicating similar Matrin3 binding patterns.

Matrin3 top targets were enriched for genes related to mitotic spindle formation and regulation and included CDC14B that is important for mitosis^42,43^. Our data demonstrate that Matrin3-dependent downregulation of CDC14B-s variant affected proliferation and phenocopied Matrin3 silencing. However, because the combination of silencing of both Matrin3 and CDC14B-s resulted in an additive effect on proliferation compared to the silencing of each gene alone, other mechanisms may be involved in Matrin3 regulation of cell proliferation.

Regarding CDC14B differential expression of variants, Matrin3 represses the inclusion of exons 13 and 14. Considering that transcription and splicing occurs simultaneously^44,45^, when Matrin3 is silenced, there is inclusion of exons 13 and 14, and exon 13 includes a premature termination codon, likely marking the transcript for degradation via NMD. In the case of CDC14B-s, Matrin3 contributes to the canonical splicing of this transcript and exclusion of exon 13 and exon 14. We decided to focus on CDC14B-s because the CDC14B-PTC is low-abundant in our RNA-seq data, consistent with it being a potential NMD target. Nevertheless, a non-coding, shorter transcript including exons 13 and 14 is found in databases (ENST00000481149), even though we do not find evidence of its existence by analysis of long-read sequencing (Iso-seq) data from HCT116 cells.

Other studies also observed that Matrin3 loss results in reduced cell proliferation^24,25,46^ and suggested increased apoptosis as a cause. Our data indicate that at least in CRC cells, Matrin3 depletion results in alterations in microtubule dynamics, possibly leading to reduced cell proliferation as studies had shown that the stabilization of microtubules inhibit cell proliferation^47–49^. Functional studies of human CDC14B protein had conflicting results regarding its role in mitosis. For instance, Berdougo et al.^36^ generated cell clones with nonfunctional CDC14B protein and showed that it did not alter the proliferation or caused spindle defects. On the other hand, Tumurbaatar et al.^43^ found that silencing of CDC14B using siRNAs reduced cell proliferation, increased the amount of bi- and multinucleated cells, and resulted in impaired metaphase-anaphase transition and caused mitotic delay that leaded to cell death. Nevertheless, other reports demonstrated that CDC14B can have an oncogenic effect through distinct mechanisms. Examples of these mechanisms include the activation of the Ras-Raf-Mek pathway, which mediates the oncogenic effect of CDC14B^50^ and also the degradation of p53, that is promoted by CDC14B phosphatase activity^51,52^. Here we suggest that CDC14B-s destabilizes microtubules and thereby causes an increase of proliferation.

Interestingly, Matrin3 may be one of several RBPs regulating microtubule dynamics while binding to the ends of polymerizing microtubules. Previously, the RNA binding properties of the classical end binding protein EB1^53^, as well as the Adenomatous Polyposis Coli (APC)^54^ scaffolding proteins were found to be required for their function, suggesting an intimate relationship between RNA regulation and microtubule dynamics.

In conclusion, we found that Matrin3 regulates splicing of CDC14B, leading to increased expression of CDC14B-s, which promotes destabilization, shorter length, and proper orientation of microtubules, that culminates in more events of mitosis and, consequently, CRC tumor growth.

## Materials and Methods

### Cell lines

HCT116, LS180, LS174T and LOVO colorectal carcinoma cell lines were purchased at American Type Culture Collection (ATCC). They were cultivated in DMEM medium (Gibco, Catalog no. 11965118) containing 10% Fetal Bovine Serum (FBS) (Gibco, Catalog no. 10082147) and 100 U/ml of penicillin and 0.1 mg/ml of streptomycin (Gibco, Catalog no. 15140122). Cells were grown at 37°C and 5% CO_2_. Cell lines were regularly checked for mycoplasma contamination using Venor™ GeM Mycoplasma Detection Kit (Sigma-Aldrich -Aldrich Co., Catalog no. MP0025-1KT).

### siRNA transfection

HCT116, LS180, LS174T and LOVO cells were reverse transfected using 20 nM siRNAs and Lipofectamine™ RNAiMAX Transfection Reagent (Invitrogen™, Catalog no. 13778075) with slight modifications from the manufacturer’s protocol. When transfections were done to purify RNA for qPCR, 1.3 to 1.75 x 10^5^ cells/per well were seeded in 12-well plates adding a mixture that was pre-incubated at room temperature for 20 minutes and contained 2 µl of lipofectamine and 1 µl of 20 µM of siRNA in 200 µl of Opti-MEM™ (Gibco, Catalog no. 31985062). The mixture was added to 800 µl of complete medium and cells.

Cells prepared for RNA-seq, protein blot or colony formation assays were reverse transfected in 6 well plates. Briefly, 3 x 10^5^ or 3.5 x 10^5^ cells/per well for HCT116 and LS180, respectively, were seeded with a solution with 5 µl of lipofectamine and 2.5 µl of 20 µM of siRNA in 500 µl of Opti-MEM™. The mixture was added to 2 ml of complete medium and cells.

Cells prepared for protein blots in Fig. 3B were transfected twice. The second round of transfection was conducted 72 hours after the first transfection and harvested 72 hours after the second transfection.

Allstars Negative Control siRNA (Qiagen, Catalog no. 1027281) were used as control siRNAs. We used SMARTPool siRNAs (Horizon Discovery, Catalog no. L-017382-00-0005) against Matrin3. In order to target CDC14B standard variant, we designed an siRNA specific for the junction of the exons 12 and 15 of it (according to the model in the Fig 3A). The abovementioned siRNA, siCDC14B-s, was purchased from Integrated DNA Technologies (IDT). For proliferation assays using those siRNAs, cells were transfected using siRNAs against more than one target (e.g. siCTRL and siMatrin3). Consequently, the final concentration of siRNAs was 40 nM. Extended Data Table 3 contains the sequences of custom siRNAs used.

### RNA extraction, RT-qPCR and RT-PCR

Cells used for expression analysis by qPCR were washed after 48 hours of transfection using DPBS 1X (Gibco, Catalog no. 14190250) and lysed using 250 or 500 µl (for 12 and 6-well plates, respectively) of TRIzol™ Reagent (Invitrogen™, Catalog no. 15596018) 48 hours post transfection. RNA was extracted according to the manufacturer’s protocol.

500 or 1000 ng of RNA was reverse transcribed using iScript™ Reverse Transcription Supermix (Biorad, Catalog no. 1708841). To the resulting 10 µl of cDNA, were added 45 or 90 µl of H_2_O, for the 500 or 1000 ng of RNA used in the reaction, respectively. 2.5 µl of diluted cDNA was used in the qPCR reaction together with 5 µl of FastStart Universal SYBR Green Master (Rox) (Millipore Sigma, Cat no. 4913914001), 0.2 µM (final concentration) of each primer and H_2_O enough for 10 µl of reaction. The reactions were done on StepOnePlus™ Real-Time PCR (Applied Biosystems™). We used GAPDH to normalize the expression and the relative expression was calculated using the 2^-ΔΔCt^ method ^55^.

Semi-quantitative RT-PCRs were made using 1µl of non-diluted cDNAs, 12.5 µl of Phusion® High-Fidelity PCR Master Mix with HF Buffer (NEB, Catalog no. M0531S) 0.8 µM of each primer (final concentration) and H_2_O enough for 25 µl of reaction. The PCR cycle used was as follows: 94°C for 5 minutes for 1 X, 98°C for 10 seconds, 50 to 60°C (CDC14B: 50°C; ST7: 56°C; CD44 and SETD5: 57°C; HP1BP3: 60°C) for 30 seconds and 72°C for 0.5 or 2 minutes (2 minutes for CD44, 1 minute for CDC14B and 0.5 minute for the rest of primers) for X28. The PCR products were resolved on 1 to 2% agarose gels.

Extended Data Table 3 contains the sequences of primers used.

### Immunostaining

For immunostaining, HCT116 cells were transfected with siRNAs as described above, using 1.5 x 10^5^ cells/per well that were seeded on coverslips (Corning, Catalog no. 2850-18) inside 6-well plates. After 48 hours of transfection, cells were fixed with ice-cold methanol for 1 minute. In case we wanted to observe cells in metaphase, we treated the cells with MG132 (Sigma Aldrich, Catalog no. 474790-10MG) at 10 µM for 90 minutes prior to fixing them.

Fixation was followed by blocking with 1% BSA in PBS/0.1% Tween (PBST) for 45 minutes at room temperature. Cells were incubated in primary antibodies for 3 hours at room temperature, washed 3 times in PBST and incubated with secondary antibodies and DAPI for 1 hour at room temperature. Following 3 washes with PBST, cells were mounted on slides using ProLong™ Gold Antifade Mounting media (Thermo Fischer, Catalog no. P36935). Mouse anti-CENP-A (Abcam, Catalog no. ab13939), rabbit anti-NUF2 (Abcam, Catalog no. ab176556), rabbit anti-Matrin3 (Bethyl Laboratories®, Catalog no. A300-591A) and mouse anti-EB1 (BD Biosciences, Catalog no. 610534) were used at 1:500 dilutions. Mouse anti-alpha tubulin (Abcam, Catalog no. ab7291) and rabbit anti-gamma tubulin (Abcam, Catalog no. ab11317) were used at 1:1000 dilutions. The secondary antibodies, goat anti-rabbit DY 488 (Thermo Fischer, Catalog no. 35552), goat anti-rabbit DY 594 (Thermo Fischer, Catalog no. 35560), goat anti-mouse DY 488 (Thermo Fischer, Catalog no. 35502), goat anti-mouse DY 594 (Thermo Fischer, Catalog no. 35510) were used at 1:500 dilution. DAPI was used at 1:5000 dilution.

### Immunoblotting

Cells used for protein blot were lysed using RIPA buffer (Life Technologies, Cat no. 89901) and sonicated for 5 seconds three times at power set of 50% (VirTis VIRSONIC 100). The lysates were centrifuged for 10 minutes at 4°C at 16 g, and the supernatant was collected. The protein was quantified using Pierce™ BCA Protein Assay Kit (Thermo Scientific, Catalog no. 23225) according to the manufacturer’s protocol. 10 to 20 µg of protein was loaded into protein (6% or 10%) SDS-polyacrylamide gels or precast gels Novex™ WedgeWell™ 4 to 20% (Thermo Fischer, Catalog no. XP04205BOX).

Proteins were transferred to a PVDF membrane using a semi-dry transfer apparatus. The membrane was blocked for 1 hour using TBST (Tris-Buffered Saline - 19.98 mM Tris, 136 mM NaCl and Tween 0.05%, pH 7.4) containing 5% milk. We used the following primary antibodies: rabbit anti-Matrin3 (Bethyl Laboratories®, Catalog no. A300-591A) at 1:2000 dilution; rabbit anti-GAPDH (Cell Signaling, Catalog no. 5174S) at 1:3000 dilution and rabbit anti-CDC14B (Thermo Fischer, Catalog no. 34-8900) at 1:500 dilution. The primary antibodies were incubated overnight, except for GAPDH (1 hour of incubation at room temperature). The membranes were developed after 1 hour of secondary antibody incubation at 1:5000 dilution by using ECL™ Prime Western Blotting Detection Reagent (Fisher Scientific, Catalog no. RPN2232).

### Immunoprecipitation

Cells prepared for immunoprecipitation (IP) were lysed as described above. 25 µl of Pierce™ Protein A/G Magnetic Beads (Thermo Fischer, Catalog no. 88803) were prepared by washing them twice with PBS 1X and RIPA buffer once. 2 µg of anti-Matrin3 or normal rabbit IgG (Cell Signaling, Catalog no. 2729S) were coupled with the beads overnight. The next day the beads were washed again with RIPA buffer and 500 µg of lysate were used for IP for 4 hours. After that, the beads were washed five times with RIPA buffer, added to SDS-PAGE sample buffer and boiled at 95°C for 5 minutes. The same was done for the input sample. Samples were loaded into a 7.5% SDS-polyacrylamide gels for Western Blot using anti-Matrin3 antibody.

### Colony formation assay

For this assay, cells were transfected with siRNAs twice. The second transfection was done48 hours after the first one. Cells were seeded after 24 hours of the second transfection. 3 x10^5^ or 3.5 x 10^5^ cells/per well were seeded in 6-well plates for HCT116 and LS180, respectively. After 8 days, cells were fixed with ice cold methanol for 15 minutes and stained with crystal violet 0.5% in methanol (10%) for 15 minutes. The ImageJ (version 2.0.0-rc-43/1.52n) software package was used to analyze images of the area of colonies.

### Proliferation assay

For proliferation assays, 1.5 x 10^3^ or 3 x 10^3^ cells/per well were seeded for HCT116 or LS180, respectively, in 96 well plates after 24 hours of transfection. Cells were incubated on Incucyte® S3 Live-Cell Analysis Instrument and photographed each 6 hours for 4 days. The pictures were analyzed by measuring the occupied area (% confluence) of cell images over time with the software from the manufacturer’s device.

### RNA-seq

RNA-Seq by poly (A) capture was performed in biological triplicates from HCT116 cells after 2 rounds of transfection of siMatrin3 or siCTRL siRNAs. The second transfection was done 72 hours after the first transfection. After 48 hours from the second transfection, cells were reseeded, and RNA was purified after 3 days from the second transfection using the RNeasy Plus Mini Kit (Qiagen, Catalog number 74134) following the manufacturer’s instructions. Samples were sequenced on HiSeq4000 using Illumina TruSeq Stranded mRNA Library Prep and paired-end sequencing.

Total RNA-seq by ribosomal RNA depletion was also performed in biological triplicates from HCT116 cells after 48 hours of siMatrin3 or siCTRL siRNAs transfection. Total RNA was extracted using TRIzol™ Reagent (Invitrogen, Catalog No. 15596026) according to manufacturer’s instructions. We used the NEBNext Ultra Directional RNA Library Prep Kit for Illumina (NEB, Catalog No. E7760) with NEBNext rRNA Depletion Kit (NEB, Catalog No. E6318). The samples were sequenced on an Illumina HiSeq 3000 machine using the 50 cycles single end sequencing protocol.

Sequence reads were aligned with the reference genome (Human - hg19) and the annotated transcripts using STAR (version 2.5.1) or TopHat (version 2.1.1) for poly (A) capture or ribosomal depletion sequencing, respectively. The gene expression quantification analysis was performed for all samples using RSEM (version 1.2.31). Differential gene expression was quantified using DESeq2 (version 1.26.0).

For transcript analysis, fastq files from poly (A) sequencing were trimmed using Trimmomatic (version 0.36)^56^ and Trim Galore (version 0.4.5) (https://www.bioinformatics.babraham.ac.uk/projects/trim_galore/). Gencode v19 version transcripts from were quantified using Salmon (version 0.14.1)^57^. The normalized TPM expression values were obtained from the Sleuth package (version 0.30.0)^58^.

### Isoform-Sequencing (Iso-Seq)

RNA from HCT116 cells was purified using the RNeasy Plus Mini Kit (Qiagen, Catalog number 74134) following the manufacturer’s instructions. The library was prepared using Iso-Seq™ Express Template Preparation protocol (Pacific Biosciences, CA, USA) for transcripts <2 kb. Raw subreads generated with PacBio SMRTlink (smrtlink-release_9.0.0.92188) were converted into HiFi circular consensus sequences (CCS). Demultiplexing was done using PacBio IsoSeq v3. The FLNC reads (Full-Length Non-Concatemer) were mapped to hg38 using Minimap2 software. Isoforms classification was done using SQANTI3.

### PAR-CLIP

Matrin-3 PAR-CLIP method was performed in three biological replicates as described previously ^26,59^. We used anti-Matrin3 antibody (Bethyl Laboratories, Catalog No. A300-591A) for the immunoprecipitation of RNPs. The resulting cDNA libraries were sequenced on an Illumina HiSeq 3000 machine as single reads with 50 cycles. Analysis was performed as described previously using PARalyzer built into the PARpipe pipeline ^33^.

### Microscopy and image analysis

Immunostained cells were imaged on Delta Vision Core system (Applied Precision / GE Healthcare, Issaquah, WA) consisting of Olympus IX70 inverted microscope (Olympus America, Inc. Melville, NY) with 100X NA 1.4 oil immersion objective and a CoolSnap HQ 12-bit camera (Photometrics, Tucson, AZ) controlled by *Softworx* software. Filters used for imaging were FITC (Ex490/20; Em 528/38), RD-TR (Ex555/28; Em 617/73) and DAPI (Ex360/40; Em 457/50) of the 86000 Sedat Quadruple Filter Set (Chroma Technology Corp, Bellows Falls, VT). Z-stacks were acquired with the thickness interval of 10 mm each. Length of mitotic spindles were analyzed by measuring distance between two poles in a mitotic spindle and length across cellular cortex to cortex aligned with the spindle poles using “measure distance” tool in *Softworx.* To prepare the figures, images were deconvolved, unless otherwise mentioned, with *Softworx* and scaled manually to 8-bit using linear LUT and the same range of scaling for all the images.

### Alternative Splicing analysis

Alternative splicing analysis was done using fastq files from RNA-Seq made by poly (A) capture using rMATs ^60^. The splicing events were filtered for the ones that appeared in at least 5 reads, in which the PSI were ≥ 0.1 or ≤ -0.1 and FDR (False Discovery Rate) <0.05 by using maser package (version 1.8.0) (http://www.bioconductor.org/packages/release/bioc/html/maser.html) for R (version 4.0.4) (https://www.R-project.org/).

### K-mer or motif analysis

We used an in-house script to make the K-mer or motif analysis. It was implemented by counting all 5-mer occurrences within the binding site sequences and comparing these values with 5-mer abundances within a background file. The background file was built by matching each binding site with random continuous sequences of equal length taken from the region of the gene where that binding site was found, like intron, 3′UTR, CDS, etc.

### Average Coverage of Binding Sites

The visualization of binding site coverage (Fig. 1H) was performed using the RCAS tool (version 1.12.0) (RCAS: an RNA centric annotation system for transcriptome-wide regions of interest) using R software (version 3.6.2).

### Statistical analysis

The statistical analysis for all data was performed using at least three replicates. The significance of statical analyses were tested using two-tailed Student’s t-test when comparing two groups or one-way ANOVA when comparing three groups. Two-way ANOVA was performed for proliferation assays. Spearman’s coefficient correlation was used for correlation analysis between different combinations of PAR-CLIP replicates samples (Scatterplots from T to C conversions). Two-tailed Kolmogorov-Smirnov test was used for comparisons of cumulative distribution. Data were considered significant when P < 0.05. Prism software (version 9) was used to make the analysis.

## Acknowledgments

This work was supported by the Intramural Research Program of the National Cancer Institute (NCI) and the National Institute for Arthritis and Musculoskeletal and Skin Disease (NIAMS). The authors would like to acknowledge Faiza Naz and Dr. Stefania Dell’Orso (NIAMS Genomics Core Facility) and Dr. Yongmei Zhao, Dr. Xiongfong Chen and Dr. Caroline Fromont (Cancer Research Technology Program, NCI) for expert help with next-generation sequencing.

## Author Contributions

A.L., M.H., and B.R.M. conceived the study. B.R.M., R.L.S., D.G.A., X.L.L., I.G. and C.C.H. developed the methodology. .B.R.M., R.L.S., D.G.A., L.P. and A.P. analyzed the data. R.L.S., D.G.A., X.L.L., I.G and M.A.B. assisted with the experiments. B.R.M., R.L.S., M.I.A., M.A.B., M.H. and A.L. wrote the manuscript. A.L. and M.H. acquired the funding. A.L. and M.H. supervised the study.

---

**Extended Data Fig 1.**
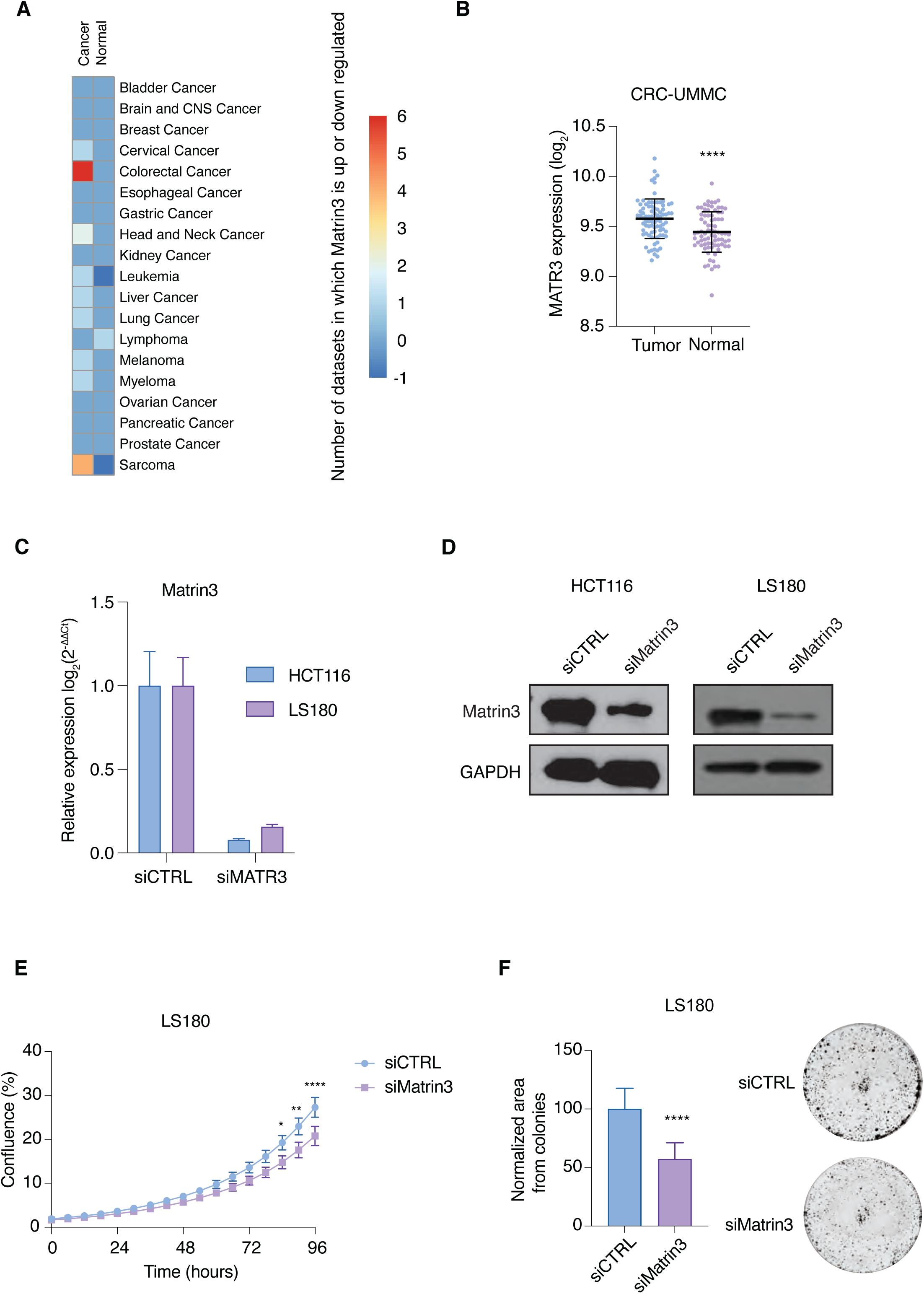
**A.** Heatmap of Matrin3 expression changes in different tumors compared to normal tissue in Oncomine datasets. Threshold was set to ≥ 2-fold expression change and p <0.0001. Matrin3 is mostly overexpressed in colorectal cancer compared to other kinds of tumors. **B.** Matrin3 RNA expression in tumors derived from colorectal cancer the from University of Maryland Medical Center (UMMC) cohort compared to respective normal samples. Unpaired, two-sided t-test; N Tumor = 83 and N Normal = 79. Error bars = SD and ****p<0.0001. **C.** RT-qPCR quantification of Matrin3 depletion using siRNAs against Matrin3 in HCT116 and LS180 cells. Error bars = SD. **D.** Immunoblots for GAPDH and Matrin3 in HCT116 and LS180 lysates following Matrin3 knockdown. **E.** Cell proliferation assay using Incucyte® after Matrin3 knockdown in LS180 cells. Two-way ANOVA test; N = 3; Error bars = SEM; F = 36.72; DF = 1, *p<0.05, **p<0.01 and ****p<0.0001. **F.** Colony formation assay after Matrin3 knockdown in LS180 cells. Unpaired, two-sided t-test; N = 3. Error bars = SD and ****p<0.0001.

**Extended Data Fig. 2.**
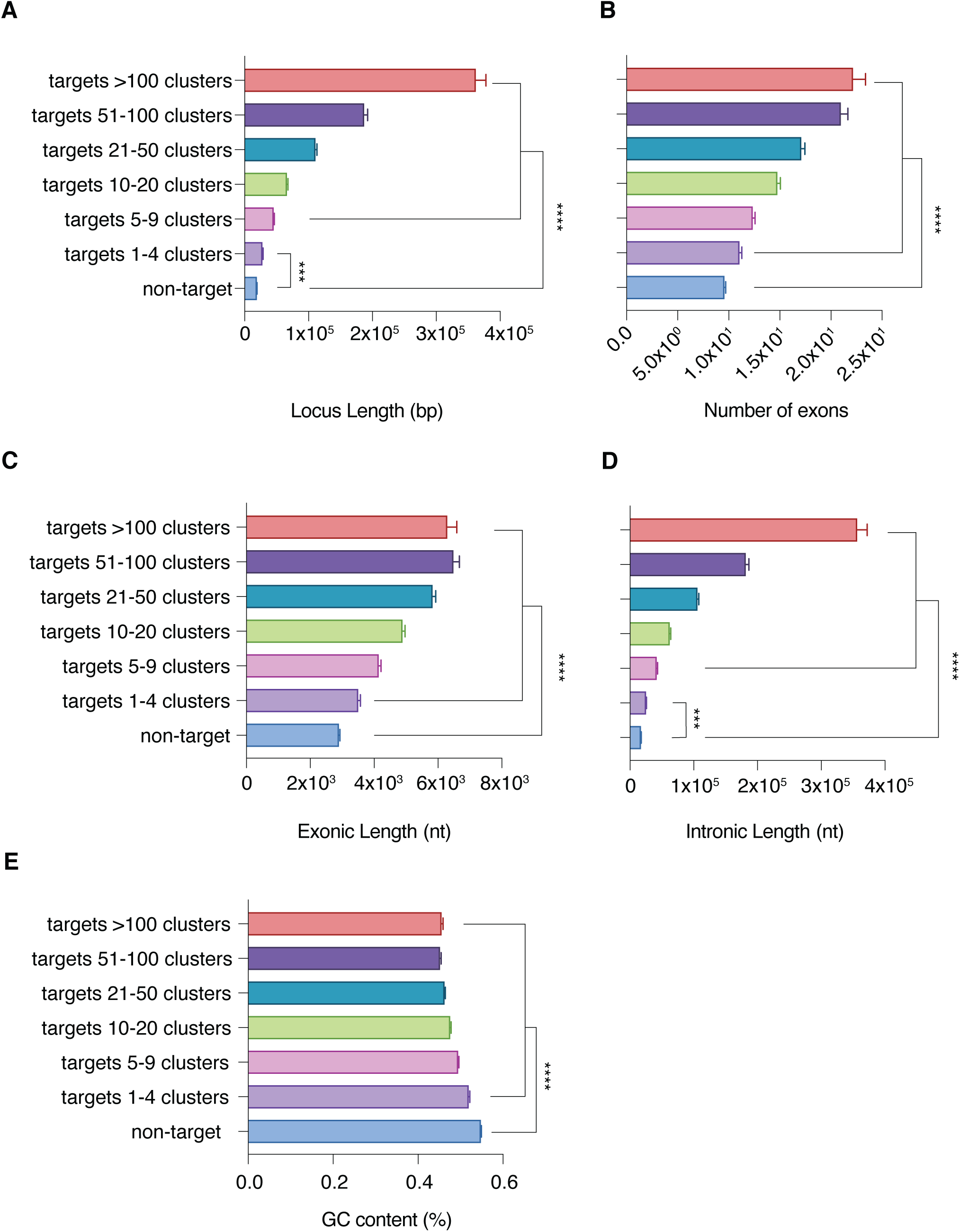
Gene architecture of Matrin3 targets. Matrin3 PAR-CLIP targets with different binding site numbers were binned according to: **A.** Locus length in base pairs. One-way ANOVA test; F = 1634; DF = 6, ***p<0.001 and ****p<0.0001. **B.** Number of exons. One-way ANOVA test; F = 192.1; DF = 6 and ****p<0.0001. **C.** Exonic length in nucleotide number (nt). One-way ANOVA test; F = 296.5; DF = 6 and ****p<0.0001. **D.** Intronic length in nucleotide number (nt). One-way ANOVA test; F = 1588, DF = 6, ***p<0.001 and ****p<0.0001. **E.** GC content in percentage (%). One-way ANOVA test; F = 322.4 and DF = 6 and ****p<0.0001. Non-targets, N = 2637; Targets with: 1-4 clusters N = 1183; 5-9 clusters N = 1185; 10-20 clusters N = 1122; 21-50 clusters N = 1007; 51-100 clusters N = 399; >100 clusters N = 167; Error bars = SEM.

**Extended Data Fig. 3.**
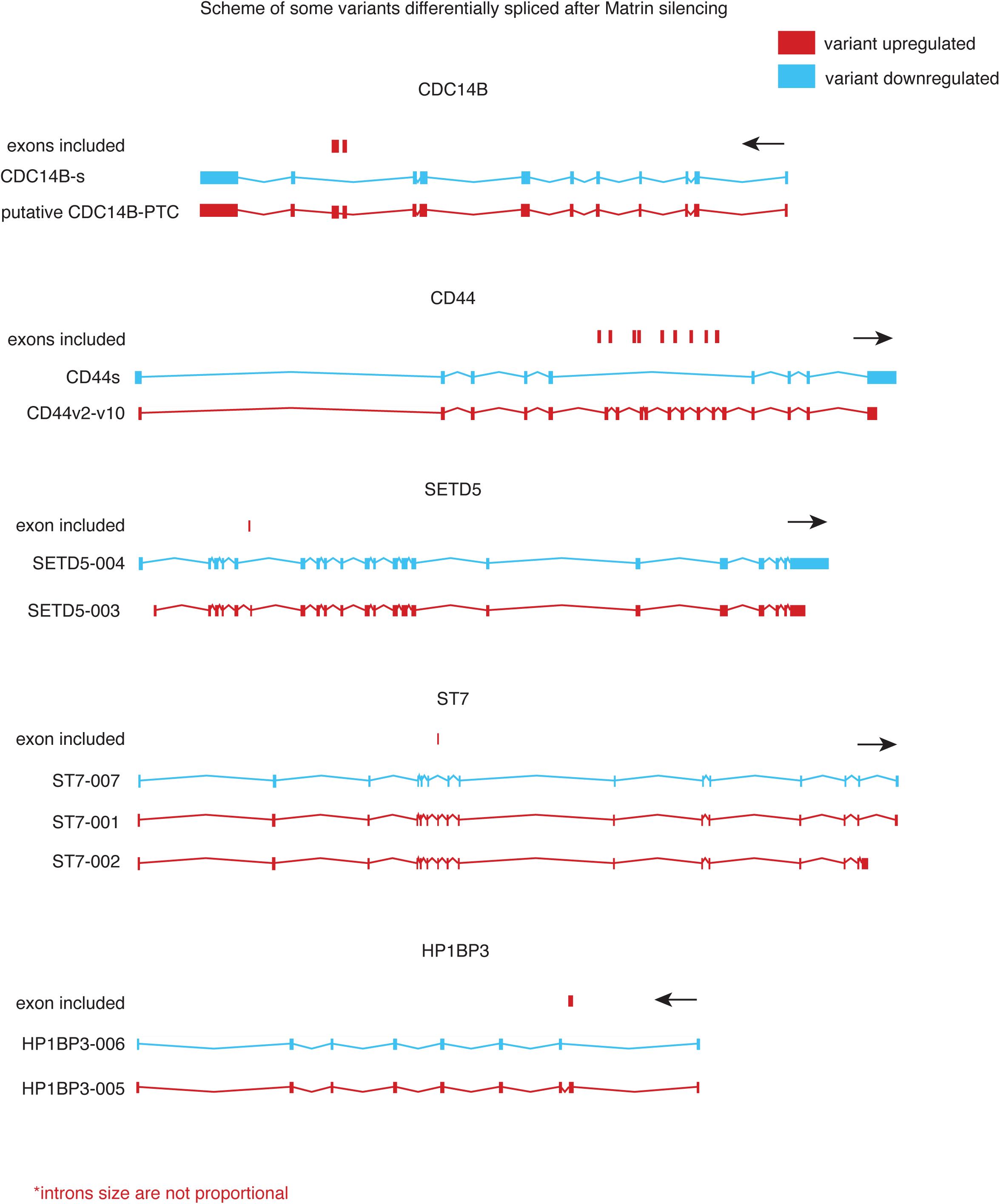
Schematic representations of variants with altered expression after Matrin3 knockdown from figures 4C-D. The image shows the exon(s) that was (were) included. The introns are not proportional to the size of the pre-mRNAs.

**Extended Data Table 1.** Number of clusters (binding sites) and crosslinked reads per gene in each Matrin3 PAR-CLIP replicate.

**Extended Data Table 2.** Number of crosslinked reads per million (NXPM or number of crosslinked reads per mRNA normalized by overall mRNA abundance) of Matrin3 RNA targets and non-targets.

**Extended Data Table 3.** Sequences of primers and custom siRNAs.

**Extended Data Table 4.** Isoforms derived from Iso-seq in HCT116 cells.

